# Computational modelling identifies key determinants of subregion-specific dopamine dynamics in the striatum

**DOI:** 10.1101/2024.07.03.601861

**Authors:** Aske L. Ejdrup, Jakob K. Dreyer, Matthew D. Lycas, Søren H. Jørgensen, Trevor W. Robbins, Jeffrey W. Dalley, Freja Herborg, Ulrik Gether

## Abstract

Striatal dopamine (DA) release regulates reward-related learning and motivation and is believed to consist of a short-lived *phasic* and continuous *tonic* component. Here, we build a large-scale three-dimensional model of extracellular DA dynamics in dorsal (DS) and ventral striatum (VS). The model predicts rapid dynamics in DS with little-to-no basal DA, and slower dynamics in the VS enabling build-up of *tonic* DA levels. These regional differences do not reflect release-related phenomena but rather differential dopamine transporter (DAT) activity. Interestingly, our simulations posit DAT nanoclustering as a possible regulator of this activity. Receptor binding simulations show that D1 receptor occupancy follows extracellular DA concentration with milliseconds delay, while D2 receptors do not respond to brief pauses in firing but rather integrate DA signal over seconds. Summarized, our model distills recent experimental observations into a computational framework that challenge prevailing paradigms of striatal DA signalling.

## Introduction

Striatal dopamine (DA) release is essential for regulating reward-related learning, incentive motivation, and motor function *(Berke, 2018; Klaus et al, 2019)*. DA exerts these roles over a broad range of time scales, yet DA primarily operates as a volume transmitter that targets metabotropic receptors located within a micrometre range from the sites of release *(Agnati et al, 1995; Borroto-Escuela et al, 2018; Cragg & Rice, 2004; Gonon et al, 2000; Sulzer et al, 2016)*. The temporal and spatial dynamics of DA release in striatum, however, remain a highly contested topic. Classically, DA release has been divided into *tonic* release, driven by pacemaker-like spontaneous firing, and *phasic* release from coordinated bursts of firing across neurons *(Niv et al, 2007; Schultz, 2007; Sulzer et al., 2016)*. However, this sharp distinction in release modes, as well as the existence of a basal DA level has recently been challenged *(Berke, 2018; Ejdrup et al, 2023; Jorgensen et al, 2023; Liu et al, 2021; Sippy & Tritsch, 2023)*.

The picture is further complicated by major regional differences across striatal sub domains. These include differences in Ca^2+^-channel and nicotinic acetylcholine receptor (nAChR) expression profiles on DA terminals, as well differential regulation and expression of the DA transporter (DAT) *(Brown et al, 2011; Cardozo & Bean, 1995; Kearney et al, 2023; Richards & Zahniser, 2009; Threlfell et al, 2010)*. In addition, we and others have found remarkable differences in extracellular DA release dynamics between the dorsal (DS) and ventral striatum (VS) *(Jorgensen et al., 2023; Mohebi et al, 2024; Salinas et al, 2023)*. Fibre photometry recordings in the DS in mice using the DA sensor dLight1.3b during self-paced exploratory activity showed a rapidly fluctuating signal, whereas we observed up to minutes-long DA dynamics in VS that correlated with behavioural output *(Jorgensen et al., 2023)*. Concurrent measurements of extracellular DA by microdialysis and fibre photometry have furthermore corroborated the lack of tonic levels of DA in DS while supported its presence in VS *(Ejdrup et al., 2023; Jorgensen et al., 2023)*. Despite these reported differences in striatal DA dynamics, electrophysiological recordings suggest that DA neurons from the primary innervators of DS, substantia nigra par compacta (SNc) and VS, ventral tegmental area (VTA), have remarkably similar firing patterns at rest *(Dodson et al, 2016)*. We therefore set out to better understand the fundamental principles governing extracellular DA dynamics by constructing a new computational model of the striatal DA system.

Extracellular DA dynamics have been modelled before; either one-dimensionally or with a primary focus on single release events or post-synaptic receptor binding *(Beyene et al, 2017; Dreyer et al, 2010; Dreyer & Hounsgaard, 2013; Dreyer et al, 2016; Venton et al, 2003; Wiencke et al, 2020)*. Here, we present a three-dimensional model of tens of thousands of release sites, focused on larger-scale signalling and based on experimentally observed biological parameters. The model faithfully replicates experimentally observed results as well as the difference in DA dynamics between DS and VS. Importantly, it offers compelling evidence that these differences do not primarily reflect different release phenomena but rather arise from differential expression and possibly nanoscale localization of the DA transporter (DAT).

## Results

### Construction of a model of DA dynamics in the striatum

We constructed a novel model of DA release using experimentally determined parameters from DS, including release, uptake, and cytoarchitecture *(Doucet et al, 1986; Dreyer et al., 2010; Dreyer & Hounsgaard, 2013; Liu et al., 2021; Olson et al, 1972; Sulzer et al., 2016)*. DA release sites on axons projecting from the midbrain were randomly simulated as uniformly distributed discrete points in a three-dimensional space (Figure 1A). The release events themselves were simulated as point source events *(Cragg & Rice, 2004)* driven by action potentials (AP). We then modelled DA release for each voxel in the simulation containing a release site as function of three key parameters; firing rate, release probability and quantal size:

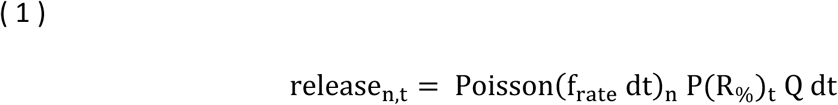

where Poisson(f_rate_ dt)_n_ is a Poisson distribution of action potentials (AP) for the given neuron (n) with the firing rate f_rate_(*1*), P(R_%_)_t_is the probability of release at the individual terminal (t) for each AP, while Q is the number of DA molecules released per event (dopaminergic quantal size) and dt the time step. Changing f_rate_ can be used to model both pacemaker firing, typically reported at 2-10 Hz, and burst firing, which can exceed 20 Hz *(Sulzer et al., 2016)*.

**Figure 1.**
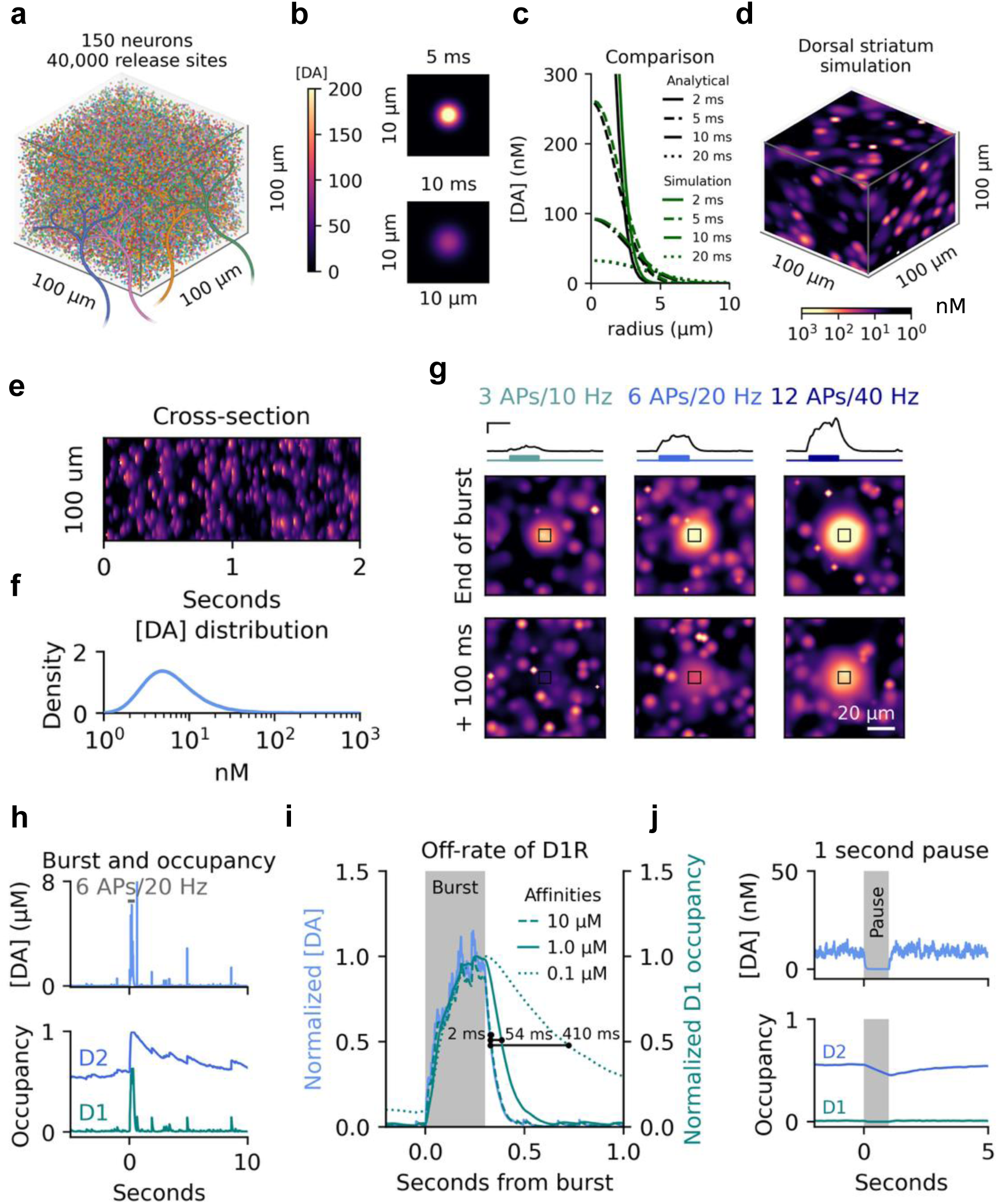
Large-scale 3D model of the dorsal striatum. (**a**) Self-enveloped simulation space of 100^3^ µm^3^ with approximately 40.000 release sites from 150 neurons. Colours of individual release sites are not matched to neurons. (**b**) Simulation of a single release event after 5 ms and 10 ms. Colour-coded by DA concentration. (**c**) Comparison of analytical solution and simulation of diffusion after a single release event at three different time points. **(d**) Representative snapshot of steady state DA dynamics at 4 Hz tonic firing with parameters mirroring the dorsal striatum. (**e**) Cross-section of temporal dynamics for a midway section through the simulation space shown in (d). (**f**) Histogram of DA concentrations ([DA]) across the entire space in (d). (**g**) DA release during three burst activity scenarios for all release sites in a 10×10×10 µm cube (black boxes) and spill-over into the surrounding space. Burst simulated as increase in firing rate on top of continued tonic firing of the surrounding space. Traces on top are average DA concentrations for the marked cubes, with bursts schematized by coloured lines below. First image row is at the end of the burst, second row another 100 ms after. Scale bars for traces are 200 ms and 500 nM. Scale bar for the images is 20 µm. (**h**) Top: representative [DA] trace for a voxel with a release site during pacemaker and burst activity. Bottom: Occupancy of D1Rs and D2Rs for same site. **(i)** Zoom on a DA burst as in (h), with [DA] in blue and D1R occupancy in teal with line style indicating different affinities. Shaded area indicates period of bursting with 6 APs at 20 Hz. (**j**) Effect of complete pause in firing for 1 second on both average [DA] and D1R and D2R occupation.

DA reuptake in the striatum is almost exclusively mediated by the DA transporter (DAT) *(Jones et al, 1998)*, which is widely distributed along DA axons and varicosities *(Block et al, 2015; Eriksen et al, 2010; Eriksen et al, 2009)*. As reuptake follows concentration-dependent Michaelis-Menten kinetics *(Nicholson, 1995)*, we simulated uptake as follows:

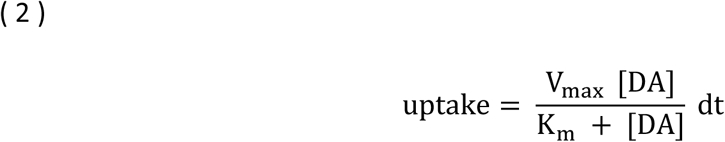

where [*DA*] is the concentration of DA for each voxel in the model, V_max_ is the maximal uptake capacity in the region and K_m_ is the concentration of DA at which half of V_max_ is reached.

The spatial distribution of released DA is a complex interplay between release, uptake and diffusion. Diffusion in an open 3D space can be simulated for each voxel with a Laplacian operator:

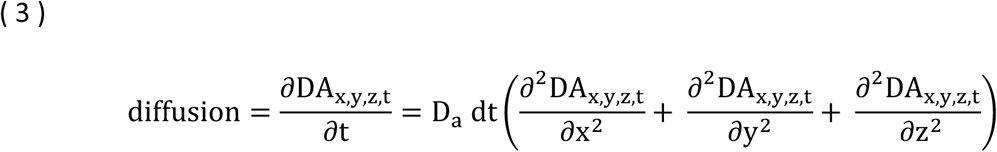

where D_a_is a corrected diffusion coefficient and *dt* is the timestep. As the extracellular space of the striatum is tortuous, we modified the conventional diffusion coefficient *D* to an apparent diffusion coefficient (D_a_) to correct for the tortuosity (λ) of the striatum *(Cragg & Rice, 2004; Nicholson, 1985)*:

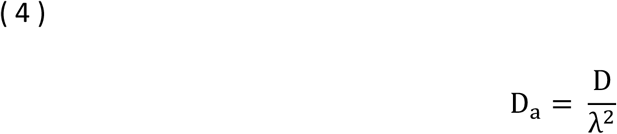

As the cerebellum exhibits as tortuosity to that recorded in the striatum, we assumed a uniform tortuosity throughout the striatum *(Nicholson & Phillips, 1981)*. Combining equation 1 through 4, we model DA changes in each voxel with a single conceptual equation:

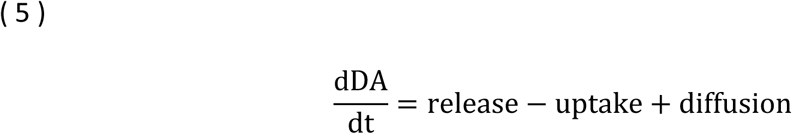

We first compared our 3D model of DA dynamics to the analytical solution of a single release event *(Cragg & Rice, 2004; Gonon et al., 2000)*. To do this, we simulated a quantal event of 3000 DA molecules and calculated DA concentrations across space at three separate time points (Figure 1B, C). The analytic solution and our model predicted almost identical results. Slight differences were introduced as the analytical solution assumes linear uptake from DAT while our model incorporate non-linear Michaelis-Menten kinetics. These differences, however, were almost negligible. The main difference between the two models lies in scalability across both space and time. Summarized, the model enables a dynamic incorporation of the surrounding DA concentration, release events and uptake, and can be scaled to cover DA dynamics of a large 3D space, whose size and granularity is only limited by computing power (see Code Availability Section for the Python code with numerical implementations of the equations listed and scripts to run the simulations and plot the main figures).

We also tested the validity of our model, by examining the response to electrical stimulation. Importantly, our model faithfully mirrored DA release seen with fast-scan cyclic voltammetry (FSCV) recordings upon direct stimulation of striatal slices when we corrected for kinetics of the typical FSCV recording setup (Figure S1b, c and discussion in Supplementary Information) *(Atcherley et al, 2015; Brimblecombe et al, 2019; Stuber et al, 2010; Xie et al, 2020)*.

### Simulating large scale DA dynamics of the dorsal striatum

To better understand the extracellular DA dynamics that arise from the balance between dopaminergic pacemaker activity and uptake, we simulated DA dynamics in the DS generated by pacemaker activity (4 Hz) of 150 neurons in the midbrain in a 100×100×100 µm space (Figure 1D, see parameters in Table 1). Our simulations yielded a pattern of partially segregated DA hot spots with large fractions of the simulated space devoid of DA, suggesting that release events in DS only elevate DA in the immediate surroundings with DAT-dependent clearance preventing a larger spread in space (Figure 1D and Video 1). This was also illustrated by a cross-section in time (Figure 1E). In line with our recent in vivo microdialysis experiments, the average DA concentration in the simulations during pacemaker activity was approximately 10 nM *(Ejdrup et al., 2023)*. Further, when we calculated the average concentration of a larger area across time, which fibre photometry conceivably does, the results resembled a tonic DA concentration (Figure S1A). However, our model predicted a spatial distribution that is highly heterogenous and devoid of pervasive resting or tonic DA levels (Figure 1D-F).

**Table 1.** List of variables used in simulation of the dorsal striatum.

| Variable | Abbreviation | Value | Reference |
| --- | --- | --- | --- |
| Firing rate |  | 4 Hz | (Paladini et al, 2003) |
| Release probability |  | 6% | (Dreyer et al., 2010) |
| DA molecules per vesicle |  | 3000 | (Klaus et al., 2019) |
| Diffusion coefficient | | $763 \mu\text{m}^2 \text{s}^{-1}$ | (Nicholson, 1995) |
| Tortuosity |  | 1.54 | (Rice & Nicholson, 1991) |
| Vmax | | $6.0 \mu\text{m s}^{-1}$ | See Table S2 |
| Km |  | 210 nM | (Hovde et al., 2019) |
| Active terminal density | - | $0.04 \mu\text{m}^{-3}$ | (Liu et al., 2021) |
| Extracellular volume fraction |  | 0.21 | (Rice & Nicholson, 1991) |
| Number of neurons in simulation space | - | 150 | (Matsuda et al., 2009) |

To ensure our simulations were performed within a sufficiently large space to yield consistent results, we tested different sizes of the simulated area and found a diameter of 50 μm to faithfully mimic the results of larger simulations (Figure S2A, B). Additionally, we tested our simulations at different granularity (0.1, 0.5, 1 and 2 μm). The finer the spatial grain, the higher detail close to a release event; however, at a spatial granularity of 1 μm, [DA] deviated by <2% across most percentiles and only by >1 nM above the 99.5^th^ percentile (Figure S2C-F), leading us to use this voxel size for our simulations.

### Burst firing and receptor occupancy

DA neurons are known to fire short bursts of APs, which is a phenomenon strongly linked to reward-prediction error and learning *(Schultz, 2007)*. These bursts can also be induced locally in the striatum by nicotinic receptor activation *(Liu et al, 2022; Matityahu et al, 2023)*.. To gain insights into extracellular DA dynamics following a locally induced burst, we simulated three different firing scenarios for a group of terminals within a 10×10×10 µm field encompassing roughly 40 release sites from the randomly simulated 150 neurons: 3 pulses at 10 Hz, 6 pulses at 20 Hz and 12 pulses at 40 Hz (Figure 1G – burst properties matched to be the same duration). The middle scenario most closely resembles the physiological burst behaviour reported in the literature, whereas the high-activity burst is above what is typically seen. Unsurprisingly, peak DA concentration was reached at the end of the bursts (Figure 1G). The 3 APs/10 Hz bursting scenario generated no significant spill-over of DA outside the region of activity, whereas the 6 APs/20 Hz and 12 APs/40 Hz bursting scenarios markedly overwhelmed uptake (Figure 1G). The relationship between firing rate and the sphere of influence by DA became further evident when plotting maximal concentration of the surrounding space (Figure S1D) and the volume of space with a DA concentration above 100 nM (Figure S1E). We found that the 3 APs/10 Hz stimulation produced DA responses that largely resembled that of a single pulse. In both cases, DA was mostly cleared after 100 ms and the volume exposed to greater than 100 nM was similar (Figure 1G and Figure S1E). In contrast, the high bursting activities caused a frequency-dependent spill-over, where the areas exposed to a DA concentration above 100 nM were 10 and 30 times larger than the terminal origin for 6 APs/20 Hz and 12 APs/40 Hz, respectively. Even after 100 ms, a considerable amount of DA remained in the 12 APs/40 Hz scenario (Figure 1G).

To understand how these DA dynamics could affect the postsynaptic response, we modelled receptor binding. D1 receptors (-Rs) were assumed to have a half maximal effective concentration (EC_50_) of 1000 nM, and we extrapolated the reverse rate constant (*k*_off_) to 19.5 s^-1^ based on a linear fit of the recently characterized DA receptor-based sensors (Figure S1E) *(Labouesse & Patriarchi, 2021)*. We set the EC_50_ of D2Rs to 7 nM and *k*_off_ to 0.2 s^-1^ based on the DA sensor kinetic fit (Figure S1F), which matches a recent binding study (0.197 s^-1^ for binding study vs. 0.204 s^-1^ based on linear fit), indicating the receptor-based sensor fit can be extrapolated to the endogenous receptors *(Agren et al, 2021)*.. To determine how these receptors would respond to our predicted DA dynamics, we simulated pacemaker activity at 4 Hz with an added burst of 6 APs/20 Hz. Figure 1H shows a representative trace of DA concentration and occupancy of the D1R and D2R for a voxel with a release site. During pacemaker activity, D1R showed an occupancy close to 0, whereas D2R occupancy was approximately 0.55 (Figure 1H). Both D1Rs and D2Rs occupancies were due to a high diffusion rate mostly invariant to individual release events caused by pacemaker activity. However, upon coordinated burst firing the occupancy rapidly increased (Figure 1H and Video 2) as diffusion no longer equilibrates the extracellular concentrations on a timescale faster than the receptors. D1R receptor occupancy closely tracked extracellular DA with a delay of only ∼50 ms for the typically reported affinity of 1 µM (Figure 1I). By contrast, it took at least 5 seconds before the burst-induced increase in D2R occupancy had declined to baseline levels (Figure 1H and Video 2). This made the D2R incapable of temporally separating closely linked bursts of activity and rather summarized the output, whereas the D1R occupancy reset between each individual burst (Figure S1G). Perhaps more surprisingly, the D2R occupancy only fell from approximately 0.55 to 0.45 when simulating a full second pause in firing due to the slow off kinetics (Figure 1J). Indeed, this finding was robust across an order of magnitude of D2R affinity (2 nm – 20 nM), although the sensitivity to a one-second pause was larger at an affinity of 20 nM (Figure S1H).

These simulations suggest that the dopaminergic architecture of the DS limits DA overflow during physiologically relevant bursting activity. Further, DA receptors had a temporally mostly uniform response to DA release caused by pacemaker activity with D1R occupancy responding rapidly to both onset and offset extracellular DA concentrations following bursts, while D2R showed seconds-long delays in offset.

### Ventral striatum maintains pervasive DA tone

Mounting evidence points to considerable differences in DA dynamics across striatal subregions *(Jorgensen et al., 2023; Mohebi et al., 2024)*, which might reflect differences in the cytoarchitectural and/or molecular dopaminergic make up. In line with this, most studies report lower dopaminergic density in the VS than in DS regardless of methodological modality with a median value of ∼90% in VS relative to DS (Table S1). Further, DAT-mediated uptake capacity is reported to be lower in VS with a median capacity at ∼30% of DS (Table S2). Consistently, we observed a clear dorsoventral gradient for DAT expression when analysing immunostainings in striatal mouse brain slices from a previous publication *(Sorensen et al, 2021)*(Figure S3A, B). By contrast, the VMAT2 staining only decreased slightly from VS to DS (Figure S3A-C).

We simulated DA release during pacemaker activity in both DS and VS. DS values were set as previously described (25 µm^3^ per terminal, uptake capacity of 6.0 μM s^-1^), but for VS we reduced the terminal density to 90% (27.8 µm^3^ per terminal) and DAT uptake capacity to 33% (2.0 µM s^-1^) (Tables S1 and S2). The remaining parameters were kept identical. With these two differences, our simulations revealed markedly different spatiotemporal DA distributions during pacemaker activity. While DS formed segregated domains with low DA concentrations in the inter-domain space (Figure 2A and Video 1), DA diffused further throughout the simulated space in VS, before being cleared by DAT. This gave rise to what may be considered a tonic DA level with hotspots of higher DA concentrations, although the concentration distribution is continuous (Figure 2A-C and Video 1).

**Figure 2.**
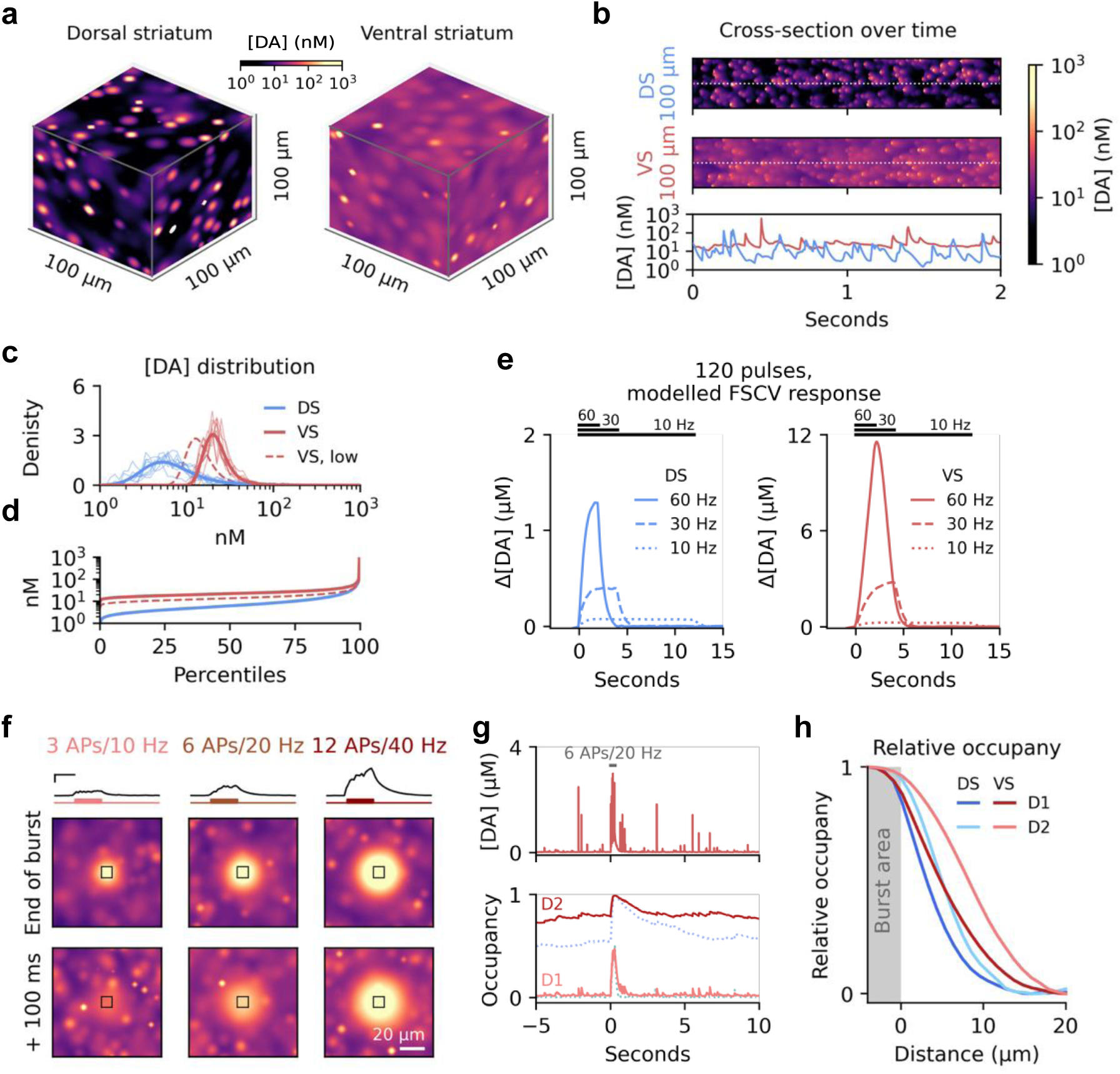
Regional difference in uptake greatly impact DA dynamics. (**a**) Representative snapshots of steady state dynamics at 4 Hz tonic firing with parameters mirroring the dorsal (left) and ventral striatum (right). (**b**) Cross-section of temporal dynamics for data shown in a. Bottom row show concentrations of the dashed lines in the top panels. (**c**) Normalized density of DA concentration of simulations in (a). Thick lines are for the entire space, thin lines are across time for five randomly sampled locations. Dashed red line is for simulation of the ventral striatum with lowest reported innervation density in the literature. (**d**) Same data as in (c), but for concentration percentiles. Note that even the lowest percentiles of VS were above 10 nM in [DA]. (**e**) Convoluted model response (Figure S1c) to mimic FSCV measurements mirroring the experimentally tested stimulation paradigm in *(May & Wightman, 1989)* for the dorsal (left) and ventral striatum (right) (**f**) DA release during three burst activity scenarios for all release sites in a 10×10×10 µm cube (black boxes) and spill-over into the surrounding space. Burst simulated as increase in firing rate on top of continued tonic firing of the surrounding space. Traces on top are average DA concentrations for the marked cubes, with bursts schematized by coloured lines below. First image row is at the end of the burst, second row another 100 ms after. Scale bars for traces are 200 ms and 500 nM. Scale bar for the images is 20 µm. (**g**) Top: representative [DA] trace 1 µm away from a release site during pacemaker and burst activity. Bottom: Occupancy of D1Rs and D2Rs for same site. Occupancy data from corresponding DS simulation on Figure 1k shown in as dotted line. (**h**) Peak occupancy at different distances from the area bursting normalized to maximal and minimum occupancy.

We compared our model of the two regions with existing experimental data. In an earlier study by May and Wightman, 120 stimulus pulses were delivered in the medial forebrain bundle (MFB) at either 10, 30 or 60 Hz and DA responses were recorded by FSCV in both caudate-putamen (CPu) and nucleus accumbens (Nac) *(May & Wightman, 1989)*. To mirror this, we simulated 120 action potentials at similar frequencies (10, 30, and 60 Hz) at 6% release probability and ran the result through convolution, as in Figure S1B and C, to generate an FSCV read-out (Figure 2E). Since May and Wightman reported no significant difference in DA released per electrically delivered pulse ([DA]_p_) between VS and DS, we applied equal quantal size and R_%_ for DS and VS in our simulations, while uptake capacity in VS was kept to a third of DS and terminal density was set to 90% as specified above.

Importantly, our simulated FSCV data closely resembled the earlier findings, with VS reaching considerably higher DA levels for all three stimulation frequencies (Figure 2E – see *(May & Wightman, 1989)*). This regional difference presumably arises from differences in DAT capacity between DS and VS, as the lower terminal density in VS would have the opposite effect (see below) and the remaining parameters were held identical.

To compare with our results for DS, we tested how VS responded during simulated burst activity. Using firing patterns identical to the DS simulations (Figure 1G), we found a larger spill-over of DA into the surrounding areas in VS (Figure 2F and Figure S3D, E). Significant amounts of extracellular DA also remained 100 ms after the physiologically relevant 6APs/20Hz firing stimulus. At the receptor level, D1R occupancy in VS showed a similar response to that in DS during the burst (Figure 1I, 2G and Video 2). By contrast, D2R occupancy during pacemaker activity was higher in VS than DS (∼0.8 versus ∼0.55 in DS), in accordance with the higher prevailing basal DA concentration. Additionally, the larger DA overflow in VS after a burst caused a higher relative increase in receptor occupancy further away from the area actively bursting than compared to DS (Figure 2H). A pause in firing had the same effect on D2R as in DS (Figure S3F).

### Changes to uptake capacity greatly affects [DA] in the ventral striatum

The values used to model the striatum (Table 1) in the previous simulations were chosen to best mimic the physiological system found in vivo. However, to test the robustness of the results, we performed simulations across wide ranges of the variable key parameters on which the model is based. First, we varied the number of varicosities actively releasing DA by setting the varicosity density to one site per 9 µm^3^ *(Doucet et al., 1986)* and simulating 4 Hz pacemaker activity with the release-capable fraction ranging from 5% to 100% (reported values range from 20% to virtually all) *(Ducrot et al, 2021; Liu et al., 2021; Liu et al, 2018; Pereira et al, 2016)* (Figure 3A). As the fraction of active sites increased, DA concentrations increased at both the median level (50^th^ percentile), which we consider a measure of tonic or baseline DA levels, and at peak levels (99.5^th^ percentile) in both DS and VS (Figure 3B, see Figure S4A for schematic of tonic and peak DA). We then used the 99.5^th^/50^th^ percentile ratio as a measure of the focality of the DA distribution (i.e. hotspot DA relative to baseline DA). This was intended as a measure of heterogeneity, that is, the higher focality, the greater competence for spatially heterogenous signalling, as has been reported in *(Hamid et al, 2021; Howe & Dombeck, 2016)*. Quantifying this across the percentage of active terminals showed that the focality of the DA distribution dropped as the active fraction increased in both regions (Figure 3C). However, the percentage of active sites in VS needed to drop to 5% to reach a relative distribution resembling the DS at a full 100% active sites, underscoring a marked difference in the spatial confinement of DA signals in VS and DS.

**Figure 3.**
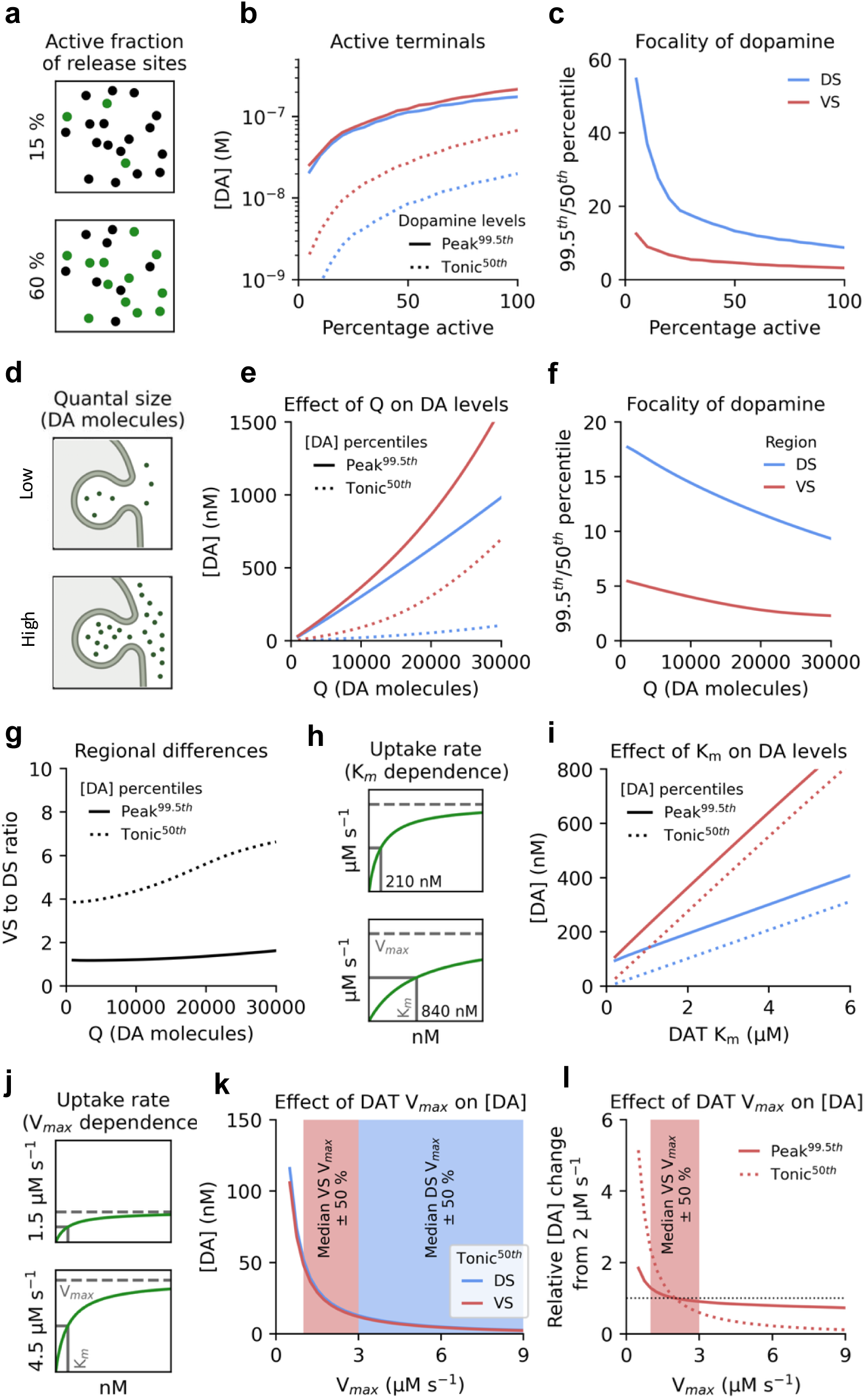
Sensitivity of the model to parameter changes. (**a**) Schematic of the fraction of active release sites. Black dots are inactive sites and green dots indicate actively releasing sites. (**b**) Effect of changing fraction of active release sites on DA concentrations. Blue line, DS peak DA concentration (99.5^th^ percentile); Red line, VS peak DA concentration (99.5^th^ percentile); Dotted blue line, DS tonic DA concentration (50^th^ percentile); Dotted red line, VS tonic DA concentration (50^th^ percentile). (**c**) Ratio between peak (99.5^th^ percentile) and tonic (50^th^ percentile) concentrations across fractions of active release sites in the DS (blue line) and VS (red line) as a measure of DA signal focality. (**d**) Schematic of changing quantal size (Q). (**e**) Effect of changing quantal size on tonic and peak DA concentrations in DS (blue lines) and VS (red lines). (**f**) Ratio between peak and tonic concentrations across various quantal sizes in in DS (blue line) and VS (red line). (**g**) Relative difference between the DS and VS for peak (black line) and tonic DA (dotted line) at different quantal sizes. (**h**) Schematic of changing DAT K_m_. (**i**) Effect of changing DAT K_m_ on DA concentrations in DS (blue lines) and VS (red lines). **j,** Schematic of changing DAT V_max_. (**k**) Effect of changing DAT V_max_ on DA concentrations. Shaded areas are median V_max_ of the two regions (DS and VS) as found in the literature shown in Table S2 ± 50%. (**l**) Effect of changing DAT V_max_, with tonic (50^th^ percentile) and peak (99.5^th^ percentile) DA concentrations normalized to their value at 2 µm s^-1^ (median value for VS). Shaded area indicates median V_max_ for VS found in the literature shown in Table S2 ± 50%.

The predicted total DA content of a vesicle and the fraction of content released per fusion event is reported to range from 1,000 to 30,000 molecules *(Garris et al, 1994; Pothos et al, 1998; Staal et al, 2004; Sulzer & Pothos, 2000)* (Figure 3D). As expected, tonic and peak concentrations increased in both DS and VS as quantal size was increased (Figure 3E). Also as expected, the focality of the DA distributions dropped for both regions as quantal size increased (Figure 3F). The relative difference in tonic DA, however, remained persistently higher in VS and even increased as quantal size increased, indicating a tendency for VS to maintain basal levels of DA regardless of release content (Figure 3G). The higher end of quantal sizes, however, resulted in median concentrations far beyond what is typically reported (Figure 3E) *(Sulzer et al., 2016)*. We observed largely similar pattern when changing either release probability or firing rate (Figure S4B-G).

DAT activity is governed by two parameters: K_m_ and V_max_ *(Kristensen et al, 2011)*. To mimic competitive inhibition of DAT by e.g., cocaine, we ran a simulation across various K_m_ values (Figure 3H) showing that increasing K_m_ caused a linear increase in DA levels, consistent with DAT uptake rate responding almost linearly to increases in [DA] below K_m_ (Figure 3I). Of note, most microdialysis studies have reported that cocaine increases [DA] to the same degree in both DS and VS; however, these quantifications are usually derived as a ratio of the absolute baseline level *(Carboni et al, 2001; Maisonneuve & Glick, 1992)*. If we divide our simulations of increasing K_m_ with the basal levels estimated in Figure 3A-C a similar response for DS and VS is found (Figure 5A). Further, we observe a convergence on a two-fold difference in the absolute values at both tonic and peak levels, which matches reports from earlier FSCV studies (Figure S5B) *(Wu et al, 2001)*. This regionally differential response to cocaine matches our observations in a previous biosensor-based study *(Jorgensen et al., 2023)*.

Finally, we changed V_max_ by ± 50% in both regions and observed a smaller change in for tonic level in DS (11 nM) than in VS (38 nM) (Figure 3K). This suggests modulation of V_max_ has higher impact in VS than DS. Further, the impact of changing V_max_ in VS was independent of both Q and R_%_ within values typically reported in the literature (Figure S5C, D). In contrast to the changes in tonic levels, the relative effect V_max_ had on peak levels was much more modest (Figure 3L).

Changes to uptake rate may be mediated by DAT internalization pathways but to our knowledge there is not in vivo evidence of differential release-uptake balances between animals that could lead to varying tonic DA levels across animals. We therefore reanalysed data from our previously published comparison of fibre photometry and microdialysis and found evidence of natural variations in the release-uptake balance of the mice (Figure S5F, G), which may underlie different tonic levels of DA in the striatum between animals.

### DAT nanoclustering affects steady state [DA] and clearance after bursts

Our simulations highlight DAT V_max_ as an effective regulator of extracellular DA levels in VS (Figure 3K). Internalization of DAT can serve as a mechanism for this control but is a relatively slow process operating on the order of minutes *(Kristensen et al., 2011)*. Interestingly, our recent studies have provided evidence that DAT move laterally in the plasma membrane and transition from a clustered to an unclustered nanoscale distribution in response to excitatory drive and other inputs *(Lycas et al, 2022; Rahbek-Clemmensen et al, 2017)*. This led us to hypothesize that DAT nanoclustering serve as a mechanism for regulating DAT activity on a faster time scale. We speculated that dense nanoclusters of DAT would produce domains of low [DA] due to uptake overpowering diffusion (Figure 4A). As uptake rate is concentration dependent, this would reduce uptake efficiency (Figure 4B). To address this hypothesis, we simulated a single ellipsoid varicosity of 1.5 μm in length and 800 nm thick with surrounding extracellular space *(Ducrot et al., 2021)*. The surface was unfolded to a square of equal area (Figure 4D), and as 9-16.4% of terminals in the striatum are estimated to be dopaminergic *(Hokfelt, 1968; Tennyson et al, 1974)*, we set the volume of the surrounding space to seven times the varicosity volume. On the surface of the varicosity, we randomly distributed eight DAT nanoclusters (Figure 4C) and ran simulations of how DAT clustering density influenced the DA clearance from the surrounding space. The observed DA concentration in the space surrounding the varicosity shown in Figure 4C is illustrated by the cross-section shown in Figure 4D. Mean DA uptake capacity of the entire space was kept constant at 4 μM s^-1^ (between the values observed for DS and VS) throughout the simulations, representing a constant amount of DAT molecules on the surface of the varicosity. We only changed the fraction of the surface of the varicosity that was uptake competent by altering the cluster size from small clusters of high density to large clusters of lower density. We ran simulations of eight identical clusters at either 20, 40, 80 or 160 nm in diameter to mirror experimentally observed cluster sizes on DA varicosities, as well as a scenario with DAT fully dispersed (Figure 4E) *(Lycas et al., 2022; Rahbek-Clemmensen et al., 2017)*. We first performed a test to see the effect clustering would have at steady state concentrations during pacemaker activity. In the unclustered scenario, average [DA] in the simulation hovered at ∼15 nM (Figure 4F). However, upon changing to a clustered architecture, [DA] rapidly increased up to 100% (∼30 nM) in just 400 ms depending on the degree of clustering (Figure 4F). We further wanted to test what effect nanoclustering would have on clearance after burst activity. To do this, we set the extracellular space to a [DA] of 100 nM and performed simulations for the same clustering scenarios (Figure 4G). Cluster size dramatically affected clearance time, with the most dense clusters taking almost 400 ms to reduce [DA] to 5 nM, compared to just ∼200 ms for the unclustered scenario. The hyper-local low-[DA] environment that arose around the dense DAT clusters became apparent when we plotted the [DA] at the centre of a cluster compared to the mean [DA] of the extracellular space (Figure 4H – unclustered also showed drop from surface of varicosity to mean of extracellular space; see general gradient of [DA] in Fig 4D). For the 20 nm cluster-scenario, the clearance was almost entirely limited by how quickly DA diffused to the nanocluster, as the local [DA] dropped to near zero. We originally hypothesized the effect would mainly be due to a depression in [DA] at the very cluster centre, but a concentration profile of the 80 nm cluster-scenario revealed the entirety of the cluster was enveloped in a low-[DA] environment (Figure 4I). Accordingly, if DA receptors are located directly next to a dense nanocluster, this effect could feasibly alter the concentration the individual receptors are exposed to.

**Figure 4.**
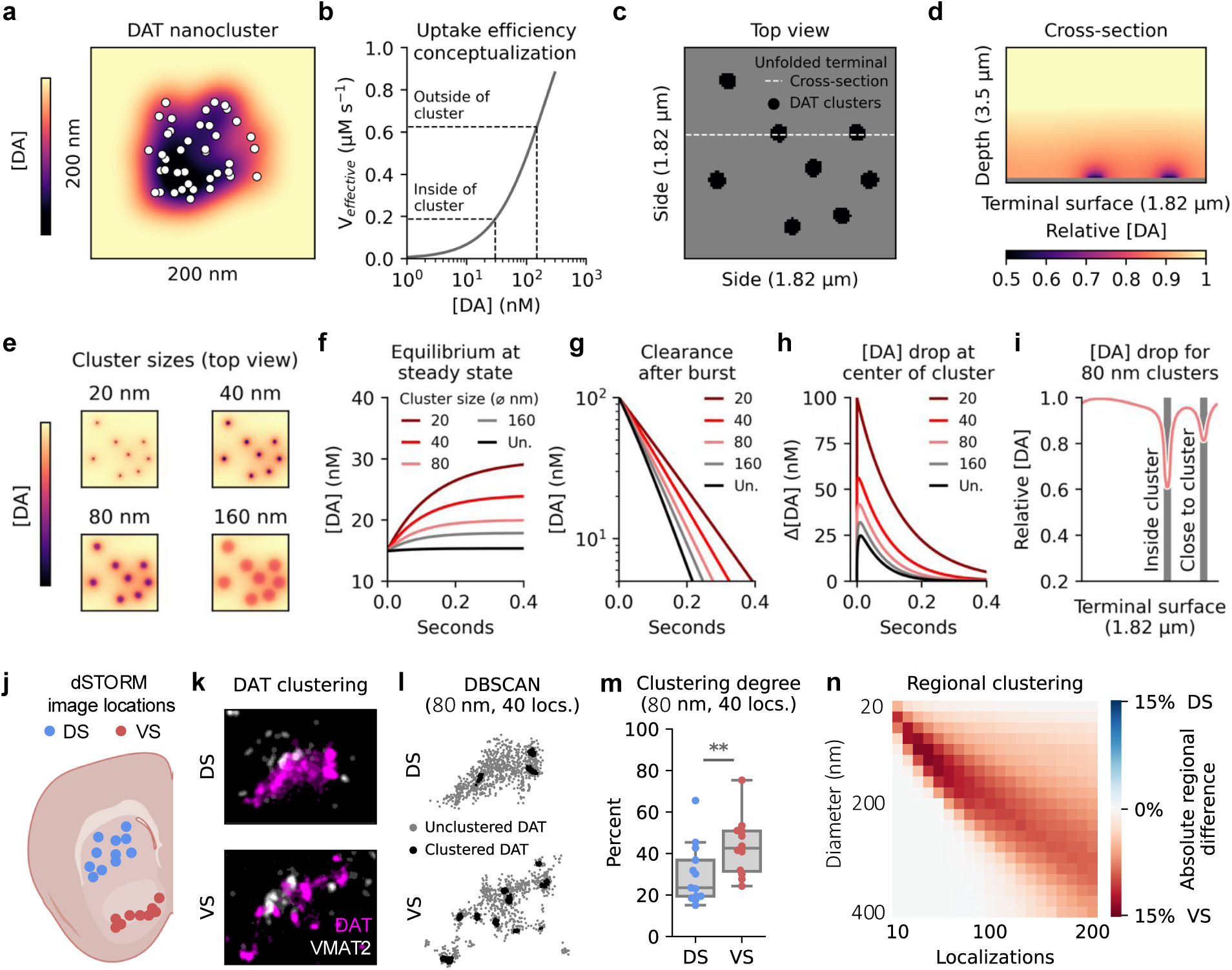
DAT nanoclustering reduces uptake and shows regional variation. (**a**) Schematic of dense DA cluster. White dots represent individual DAT molecules, and colour gradient the surrounding DA concentration. (**b**) Effective transport rate dependent on local concentration. (**c**) Top view of unfolded DA varicosity. Black shapes denote clusters of DAT. Dashed white line indicates placement of cross-section shown in (d). (**d**) Cross section showing DA concentration in space surrounding varicosity unfolded in c. Grey line at the bottom is surface of the varicosity. Colour-coded for DA concentration. (**e**) Top view from (c), but colour coded for DA concentration immediately above membrane surface at different DAT cluster sizes. (**f**) Changes in [DA] from 15 nM unclustered (Un.) steady state with constant release after changing to four different cluster size scenarios (ø = diameter). (**g**) Clearance of 100 nM [DA] for different DAT cluster sizes. (**h**) Difference between DA concentration at the centre of clusters (or general surface of varicosity for unclustered) and mean concentration of the full simulation space (**i**) Concentrations across a cross section of a surface with 80 nm diameter clusters. Shaded areas highlight cluster locations. (**j**) Location of images of the dorsal (DS) and ventral striatum (VS) in striatal slices from mice as imaged in *(Sorensen et al., 2021)* with direct stochastic optical reconstruction microscopy (dSTORM). (**k**) Two representative DA varicosities from DS and VS with VMAT2 in white and DAT in magenta. Images are 1.5×2 µm. (**l**) Individual DAT localizations (locs.) from images in (k) coloured by clustering. Black indicates localization identified as clustered based on DBSCAN with parameters 80 nm diameter and 40 localizations. Grey indicates unclustered localizations. (**m**) Quantification of clustering across all images in (j) with parameters in (l). Welch’s two-sample t-test, P = 0.012(*), n = 12 (DS) and 13 (VS). (**n**) Absolute difference in percentage of clustering as assessed with DBSCAN across a range of parameters. VS has a higher propensity to cluster across cluster sizes typically reported for DAT clusters.

Summarized, the data supports that DAT nanoclusters produce domains of low [DA] as uptake outcompetes diffusion. As transporter uptake is concentration dependent, the overall uptake efficiency is reduced, which in turn may lead to higher extracellular concentrations of DA. The simulations therefore posit DAT nanoclustering as an efficient way to regulate the extracellular levels of both tonic DA and the spatiotemporal [DA] profiles following release.

### DAT clusters more in the ventral striatum

As our simulations suggest that nanodomain clustering is a way of regulating uptake, and this may be particularly efficient at controlling extracellular DA in VS, we hypothesized nanoscale clustering would be more prominent in VS. To investigate this, we reanalysed data from a previous publication *(Sorensen et al., 2021)*, where we acquired super-resolution images of coronally sliced striatal sections from mice stained for DAT. For the present study, images were split in DS and VS based on where in the slices they were taken (Figure 4J). DA terminals were identified by vesicular monoamine transporter 2 (VMAT2) expression, and qualitatively DAT appeared more clustered in VS than DS (Figure 4K). To quantify this, we applied the clustering algorithm Density-Based Spatial Clustering of Applications with Noise (DBSCAN) (Figure 4L). This confirmed a regional difference with more DAT localized clusters when using DBSCAN to identify clusters with scanning diameter of 80 nm in VS compared to DS (Figure 4L, M), and we observed a similar regional difference across the range of cluster sizes typically reported (20 to 200 nm; Figure 4N) *(Lycas et al., 2022; Rahbek-Clemmensen et al., 2017)* supporting the conclusion that DAT nanoclustering is more prevalent in VS than DS.

## Discussion

We developed a three-dimensional, finite-difference computational model to investigate spatiotemporal DA dynamics of striatal subregions in detail. Leveraging prior experimental information on regional differences in dopaminergic innervation density and DAT uptake capacity, our model predicts important differences in dopaminergic dynamics between DS and VS. Strikingly, our simulations suggest that large areas of the DS are effectively devoid of a basal level of DA at pacemaker activity, whereas VS maintain a more homogenous tonic-like basal DA concentration with only small changes in uptake activity powerfully regulating the extracellular DA tone. Furthermore, we modelled receptor binding kinetics and found that D1R binding faithfully followed recently described rapid DA dynamics of the striatum *(Ejdrup et al., 2023; Jorgensen et al., 2023; Markowitz et al, 2023)*, while D2R, with an off-rate of ∼5 seconds, appeared better suited for detecting background tone and integrating prolonged activity *(Howe et al, 2013; Jorgensen et al., 2023)*. Collectively, these observations have important implications for our understanding of striatal function in behaviour, including decoding of inputs from the prefrontal cortex (PFC) and limbic system as well as the influential phasic-tonic model of dopaminergic signalling *(Grace et al, 2007; Niv et al., 2007; Schultz, 2007)*

It has been assumed for long that there are tonic levels of DA in the striatum *(Niv et al., 2007; Schultz, 2007; Sulzer et al., 2016)*, although the phenomenon has no clear definition *(Berke, 2018)*. Our simulation of DA dynamics in DS during pacemaker activity showed no evidence for a homogenous extracellular distribution. Rather, elevated [DA] was transiently present around release sites during pacemaker activity with the remaining space mostly depleted of DA. The absence of a general tonic DA level in DS predicted by our model directly supports the notion that DA release sites in DS establish distinct and only partially overlapping DA domains rather than diffuse, tonic DA levels (see *(Liu et al., 2021)*). This conclusion aligns with recent data where we found that DA concentrations measured by microdialysis correlates with the average of rapid activity recorded with fibre photometry rather than a baseline DA tone *(Ejdrup et al., 2023)*. Earlier modelling work by Wickens and colleagues predicted pacemaker activity would generate a tonic, uniform concentration *(Arbuthnott & Wickens, 2007)*. But our modelling suggested this is prevented by the significant DA uptake capacity of the DS, as measured by more recent reuptake studies (see Table S2).

In contrast to the DS, our model predicted VS to hold a considerable basal level of DA even in spaces without an immediately adjacent release site. This is conceivably what most refer to as tonic DA. In this study, we quantified tonic DA as the median concentration of the entire space (50^th^ percentile), which appeared significantly higher in VS than DS because of the lower VS uptake capacity. Importantly, this matches the results of our direct in vivo comparison of DS and VS in freely moving mice *(Jorgensen et al., 2023)*, as well as supporting a spatial gradient of time horizons in the striatum previously predicted in a separate work by Wickens and colleagues *(Wickens et al, 2007)* and measured in vivo by Mohebi and colleagues *(Mohebi et al., 2024)*.

To challenge our model predictions, we performed our simulations across a wide range of parameters. Only changes to V_max_ for uptake generated differential responses in the two regions. With release and uptake parameters at values from the literature, VS was at a critical point where minor changes to uptake significantly impacted the tonic levels without any major effect on peak concentrations.

Contrary to our observations, some previous microdialysis experiments have suggested higher basal DA levels in the DS compared to VS (Kuczenski & Segal, 1992; Shen et al., 2004 have reported 2-4 times higher [DA] in DS compared to VS *(Kuczenski & Segal, 1992; Shen et al, 2004)*.However, there are disparate observations in the literature (e.g. Carboni et al., 2001 reported 20% higher [DA] in VS *(Carboni et al., 2001)*. Moreover, it is important to note that regional comparisons in microdialysis might be confounded by the considerably higher uptake rate in DS. This will increase the extraction fraction and possibly lead to a significant overestimation of the extracellular concentration as compared to a region with lower uptake rate, such as the VS *(Chefer et al, 2009)*.

Our three-dimensional simulations highlight DAT-mediated reuptake as a key mechanism governing striatal DA dynamics and as a key mediator of regional-specific DA dynamics. A physiologically relevant way to regulate uptake capacity is moving DAT to-and-from the plasma membrane. Indeed, DAT is subject to such regulation and some of these mechanisms may even be exclusive to the ventral region, including protein kinase C-induced DAT internalization and Vav2 regulation of DAT surface expression *(Fagan et al, 2020; Zhu et al, 2015)*. Chemogenetic G_q_-coupled DREADD activation of DA neurons also result in differential DAT trafficking in the two regions *(Fagan et al., 2020; Kearney et al., 2023)*. The findings position DAT regulation as an excellent candidate for changing tonic DA levels in VS, which has been proposed to selectively attenuate afferent drive from the PFC through D2R activation *(Grace et al., 2007)*. Recent studies of D2R expressing spiny projection neurons (SPNs) in the VS also suggest that the receptor is not fully saturated under basal firing *(Lee et al, 2020)*, matching both our simulations of receptor binding and the notion that tonic DA can be manipulated to alter D2R activation.

If changing uptake capacity is to have a behavioural relevance on a fast timescale, a mechanism to regulate DAT function faster than internalization must exist. Importantly, the transporter does not only move to-and-from the surface, but also laterally in the plasma membrane. We have reported that DAT forms nanoclusters in the plasma membrane that dynamically reshape based on excitatory and inhibitory input *(Lycas et al., 2022; Rahbek-Clemmensen et al., 2017)*. Moreover, we have previously shown that cocaine, which both competitively inhibits DAT and reorganizes the transporter nanodomains *(Lycas et al., 2022)*, changes the DA signal of the DS to dynamics akin to the VS *(Jorgensen et al., 2023)*. Importantly, our simulations showed that nanoclustering may be an effective way to sequester DAT in a dense domain where uptake overpowers diffusion and as a result brings down effective uptake speed through local DA depletion. This is in line with evidence that these DAT nanoclusters are enriched in phosphatidylinositol-4,5-bisphosphate (PIP2), and that metabolism of PIP2 decreases uptake rate of DAT *(Lycas et al., 2022)*; Carvelli, 2002 #2339}. We also found that the nanoclustering phenomenon was considerably more prevalent in VS than in DS. Taken together, these data point to DAT nanoclustering as a way to shape both the spatiotemporal profile of DA release as well as the tonic levels of DA in the striatum – particularly in the VS.

Our incorporation of receptor binding was inspired by important previous modelling work *(Dreyer et al., 2010; Dreyer & Hounsgaard, 2013; Hunger et al, 2020)*. However, earlier models by Dreyer & colleagues assumed instantaneous equilibrium between extracellular DA and receptor occupation, which disregards differences in kinetics of the DA receptors that greatly impact transmission dynamics. While later work by Hunger & colleagues introduce more complex receptor modelling, they based their kinetics parameters on early pharmacological studies, whose values likely would prevent DA receptors from decoding signal below the order of minutes *(Burt et al, 1976; Maeno, 1982; Nishikori et al, 1980; Sano et al, 1979)*. Instead, we based our receptor kinetics on newer pharmacological experiments in live cells *(Agren et al., 2021)* and properties of the recently developed DA receptor-based biosensors *(Labouesse & Patriarchi, 2021)*, whose receptor values match well despite different methodological approaches. The biosensors are mutated receptors whose kinetics may not be identical to the endogenous receptors, but only the intracellular domains are altered, with no apparent changes of the binding site *(Labouesse & Patriarchi, 2021)*.

Indeed, these biosensors exhibit kinetics that are well aligned with both modelled and experimentally reported extracellular DA dynamics using non-biosensor-based methods *(Atcherley et al., 2015; Gonon et al., 2000; Venton et al, 2002)*. We believe accordingly that our updated parameters are more accurate portrayals of *in vivo* conditions; however, as shown throughout the study, the affinity values greatly affect the results. Therefore, we find it important that our model will be available to the research community, allowing others to test their own estimates of receptor kinetics and assess their impact on the model’s behaviour.

The presented simulations suggested that receptor binding was largely invariant to single release events during pacemaker activity, while bursts of activity rapidly changed occupancy. Both D1R and D2R immediately responded to burst onset; however, while D1R occupancy rapidly declined to zero within approximately 50 ms, the slow D2R kinetics resulted in an occupancy decline over ∼5 seconds, returning to the baseline maintained by tonic firing. This means that D1R is better suited to discriminate inputs in rapid succession and allow for postsynaptic decoding of the fast-paced in vivo dynamics described in particularly for DS *(Ejdrup et al., 2023; Jorgensen et al., 2023; Markowitz et al., 2023)*. By contrast, our analysis shows that D2Rs integrate DA signals over several seconds. As D1R occupancy is negligible during pacemaker activity and D2R kinetics are too slow to pick up rapid changes in DA concentration, our simulations moreover suggest pauses in firing of less than 1 second is not an effective way of signalling for the striatal dopaminergic system.

Notably, this finding was apparent even when the D2 affinity was increased an order of magnitude. This challenges the effectiveness of proposed negative reward prediction errors, as even a long pause in firing would have a limited effect on D2 receptor occupation and downstream signaling. This may explain why DA drops during reward omissions are not nearly as prominent as positive signals *(Farrell et al, 2022; Greenstreet et al, 2025)*..

In conclusion, we have developed a three-dimensional model for DA release dynamics and receptor binding that integrates a wealth of experimentally determined parameters and generates responses to electrical and pharmacological input that fits robustly with literature observations. The model offers an important theoretical framework and a predictive tool that can serve as the basis for future experimental endeavours and help guide the interpretation of new as well as older empirical findings on DA signalling dynamics under both physiological conditions and in disease.

## Methods

### Three-dimensional finite difference model

DA release sites in the striatum were simulated at a density of one site per 25 µm^3^ in DS and one site per 27.8 µm^3^ in VS (except for Figure 4b and c). As a single medium spiny neuron in the striatum is estimated to be influenced by axons from 95 to 194 dopaminergic neurons *(Matsuda et al, 2009)*, we simulated DA release sites as originating from 150 separate cell bodies. These were set to fire independently from each other at 4 Hz generated by a Poisson distribution. We set the release probability in response to an action potential at the individual terminal levels to 6% *(Dreyer et al., 2010)*, and if a release occurred the DA concentration of the voxel was elevated by the equivalent of 3000 DA molecules as per Equation 1. The volume of each voxel was corrected for an extracellular volume fraction of 0.21 *(Rice & Nicholson, 1991)*.

We did not incorporate paired-pulse depression in our model as most in vivo FSCV studies have not reported this phenomenon *(Chergui et al, 1994; Garris & Wightman, 1994; Hamid et al, 2016; May & Wightman, 1989)*. In addition, we could explain the gradual blunt of the response during prolonged stimulation (Figure 2E) as the result of an equilibrium between release and uptake rather than a depression in release. Newer in vivo biosensor-based studies observe the same “blunting” phenomenon during stimulation trains *(Mohebi et al, 2023)*. A low initial release probability that scales well with frequency has also been reported for VMAT2-positive release sites in dopaminergic cultures where the axons are not severed from their cell bodies as in acute striatal slices *(Silm et al, 2019)*. Note that we did not explicitly model autoreceptor inhibition at pacemaker activity. However, this value should be implicitly reflected in the in vivo estimates we base our model on.

When simulating bursts or trains of stimulations, action potentials were added as described for each simulation, but release probability was kept the same unless otherwise noted. Uptake was evenly distributed throughout the simulation space with a V_max_ at 6 µm s^-1^ for DS and 2 µm s^-1^ for VS (see Table S2) and a K_m_ at 210 nM *(Hovde et al, 2019)* and calculated for each voxel as per Equation 2. For each time step, *dt*, diffusion calculated with a finite difference implementation of the Laplacian operator described in Equation 3 using a diffusion coefficient corrected for tortuosity at 321.7 µm^2^ s^-1^. Periodic boundaries were used to avoid edge issues with a simulation size of 50×50×50 µm unless otherwise noted. The Python-based implementation of the model can be found as described in the Code and Data Availability section.

### Receptor occupancy

Our modelling of receptor binding was based on the work of (*Dreyer & Hounsgaard, 2013).* Occupancy of the receptors for any given voxel was calculated numerically by numerical integration of:

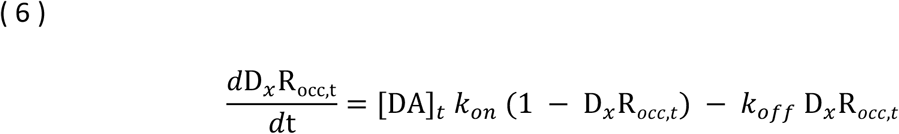

Where [DA]_t_ was the dopamine concentration at the given voxel, D_x_R_occ,t_ the fraction of DA recptors occupied in the same voxel, *k*_on_ the rate constant and *k*_off_ the reverse rate constant. The model assumed no buffering effect from DA bound to receptors, as DA receptors are GPCRs typically expressed at very low levels *(Zhang et al, 2024)*.

### Immunohistochemistry analysis

Analysed widefield images were obtained from *(Sorensen et al., 2021)*. The mouse striatum slices were stained for either DAT or VMAT2 (see Sørensen et al., 2021 for slicing, fixation, staining and imaging protocol *(Sorensen et al., 2021)*. Fluorescence intensities were extracted in a single dorsoventral line parallel to the midline through the horizontally centre of the anterior commissure using ImageJ v1.53q. for each slice. Data was vertically centred around the anterior commissure. No background subtraction was performed.

### FSCV modelling

FSCV recordings were modelled by convoluting a simulated DA response to a pseudo FSCV trace. The impulse function used was as described in *(Venton et al., 2002)*. In brief, the impulse function was defined as IF_FSCV_ = e^-(t+1)(k^_-1_^*t^_s_^+k^_-2_^*t^_o_^)^, where t is time, t_>_is scan time, t_?_ is oxidization time, and k_7:_ and k_7-_ desorption constants. t_>_ was set to 100 ms as reported in *(May & Wightman, 1989)* and t_?_ to 4 ms, which is half of the typically reported symmetrical waveform length. The desorption constants (k_-1_ = 1.2 s^-1^ and k_-2_ = 12 s^-1^) were taken directly from *(Venton et al., 2002)*.

### Nanocluster uptake modelling

Nanoclustering of DAT was simulated in Python. An ellipsoid-shaped varicosity of 1.5 μm in length and 800 nm in width was modelled as a flat, square surface with a width of 1.8 μm to match the total surface area *(Lycas et al., 2022; Rahbek-Clemmensen et al., 2017)*. As 9-16.4% of terminals in the striatum are estimated to be dopaminergic*(Hokfelt, 1968; Tennyson et al., 1974)*, we set the height of the simulation space to 3.52 μm to generate an extracellular volume seven times the size of the varicosity. Axons and cell bodies are likely a non-negligible part of the striatal volume but omitted in this analysis due to a lack of information in the literature (however, decreasing the share of striatal space taken up by DA varicosities works in the direction of our hypothesis). Spatial granularity was set to 20 nm, and we randomly placed eight non-overlapping locations for clusters on the simulated varicosity. Uptake capacity per voxel for each clustering scenario was set so the mean uptake of the entire space was 4 μm s^-1^, which resulted in clusters with increased DAT density with decreasing cluster size.

For the steady state simulations, we added [DA] at a constant rate of 224 nM s^-1^ calculated as follows:

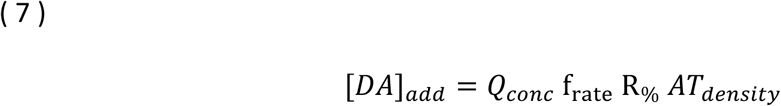

Where [*DA*]*_add_* was the DA concentration added per second, *Q_conc_* the DA concentration added to a voxel by a single release event, f_rate_ the firing rate at pacemaker activity, R_%_ the probability of release for each action potential and *AT_density_* the density of actively releasing terminals. *Q_conc_* was calculated as follows:

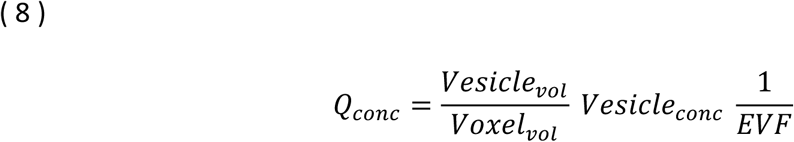

Where *Vesicle_vol_* is the volume of a vesicle (with an assumed radius of 25 nm), *Voxel_vol_* the volume of a voxel in the simulation space, *Vesicle_conc_* the concentration of DA in a single vesicle (assuming 3000 DA molecules) and *EVF* the extracellular volume fraction of the striatum.

For the clearance after burst simulations, we set the initial DA concentration of the entire space to 100 nM.

We ran all the nanoclustering simulations with a time step of 1.875e^-7^ s to avoid numerical instability. Diffusion corrected for tortuosity was set to 321.7 µm^2^ s^-1^ and calculated with a Laplacian operation as described in Equation 3. Periodic boundaries were used to account for the varicosity being circular and avoid issues at the edge of the simulation space. The full implementation in Python can be found as described in the Code and Data Availability section.

### Super-resolution clustering analysis

Super-resolved fluorescence microscopy dual color images from *(Sorensen et al., 2021)* of striatal brain slices from mice were analyzed based on their position in the dorsal or ventral regions (Figure 4J). Localizations were fitted using DAOSTORM *(Babcock et al, 2012)* on the originally acquired direct stochastic optical reconstruction microscopy (dSTORM) videos with the following settings: background sigma of 8, maximum likelihood estimation as the fitting error model, 20 peak identification iterations, an initial sigma estimate of 1.5 and a threshold adjusted to each imaging session. Each image was analysed for percentage of localizations within a cluster using density-based spatial clustering of applications with noise (DBSCAN) *(Ester et al, 1996)* using the Python package Scikit-learn 0.22 *(Pedregosa et al, 2021)*. To uncover broader trends, we scanned across a range of parameters: radius was run from 10 nm to 200 nm and number of localizations from 10 to 200. For Figure 4M a diameter of 80 nm was chosen to match the size reported in *(Rahbek-Clemmensen et al., 2017)*.

## Statistical Analysis

The statistical analyses performed can be found in the legends of each figure. Statistical analyses were performed using open-source python packages SciPy v1.5.2 and NumPy v1.18.1. Boxplots show the 25th and 75th percentile range and whiskers indicate up to 1.5 times the interquartile range. The remaining data points are plotted as individual outliers.

## Acknowledgements

We thank Dr. Kenneth L. Madsen, University of Copenhagen for general input on our modelling and Dr. Nicolas Tritsch, New York University Grossman School of Medicine, for a fruitful discussion on the postsynaptic receptor properties and occupation.

## Additional Information

### Competing Interest

Authors declare that they have no competing interests.

### Funding

The work was supported by the Lundbeck Foundation grants R266-2017-4331 (UG), R276-2018-792 (UG), R359-2020-2301 (UG), R181-2014-3090 (FH) and R303-2018-3540 (FH).

### Author Contributions

ALE, FH and UG conceptualized the study. ALE built the model with help from JKD. ALE ran the simulations. MDL and SHJ provided technical input on experimental data. ALE drafted the manuscript. ALE, FH and UG wrote the final version of the manuscript with comments and suggestions from JKD, TWR and JWD. All authors critically evaluated the manuscript. FH and UG supervised the project.

## Additional files

### Supplementary file

Supplementary Text.

Figures S1 to S5.

Tables S1 and S2.

Videos 1 and 2.

### Data Availability

Numerical implementation and code to run the simulations are available at: https://github.com/GetherLab/striatal-dopamine-modelling. All experimental data are from *(Ejdrup et al., 2023)* or *(Sorensen et al., 2021)*. Corresponding authors on these papers should be consulted for data availability.

## Notes

https://github.com/GetherLab/striatal-dopamine-modelling

### Competing Interest Statement

JKD was employed at H. Lundbeck A/S while conducting this study.

### Summary of Updates

The revisions herein include both several new and modified simulations based on highly constructive feedback from reviewers: – Comparison of analytical and simulation of diffusion after just 2 ms (Figure 1B) – D1 off-rates at various affinities (Figure 1I) – Equilibrium of DA levels at various DAT nanocluster sizes (Figure 4F) – D1 and D2 occupancy after chain of bursts (Figure S1G) – D2 occupancy in response to pause in firing at various affinities (Figure S1H) We believe that these new simulations, together with a thorough revision of the text, have both further substantiated the biological significance of our manuscript and further supported our conclusions.

