## Supplemental information for "Computational modelling identifies key determinants of subregion-specific dopamine dynamics in the striatum"

### Supplementary Text

#### Burst release and FSCV modelling

To test the validity of our model, we examined how it would respond to a simulated direct electrical stimulation. We chose this experimental paradigm, as direct stimulation of striatal slices with fast-scan cyclic voltammetry (FSCV) recordings of DA is the most common procedure in the literature to measure DA release. However, only around 10% of the DA release in such a procedure stems from direct stimulation of DA terminals as 90% can be attributed to cholinergic release from interneurons activating presynaptic nicotinic receptors on the pre-synapse of dopaminergic axons *(Liu et al, 2022; Liu et al, 2018; Zhou et al, 2001)*. This leads to a much stronger response than an action potential travelling up from the midbrain would generate, which is estimated to have an *in vivo* release probability on the order of 6% *(Dreyer et al, 2010)*. We therefore simulated an event with 10x release probability (60%) to scale to the expected output from an electrical stimulation in slices. The numbers for quantal size vary throughout the literature, but for consistency with previous simulations we set the value to 3000 DA molecules *(Cragg & Rice, 2004; Omiatek et al, 2010; Pothos et al, 1998; Sulzer & Pothos, 2000)*. On the uptake side, DAT has a $V_{max}$ in DS of roughly 6 μM $s^{-1}$ and a $K_m$ of 210 nM (Table S2) *(Calipari et al, 2012; Dreyer et al., 2010; Hovde et al, 2019; May & Wightman, 1989; Salinas et al, 2023; Siciliano et al, 2014)*. In response to such a stimulation, mean DA concentration ([DA]) at an artificial carbon probe immediately rose to 200 nM with full return to baseline in roughly 0.2 seconds (Extended Data Fig. 1B, C). This is significantly faster dynamics than what is typically observed using FSCV, but similar to newer observations made in the DS measured with the dLight biosensors *(Atcherley et al, 2015; Brimblecombe et al, 2019; Mohebi et al, 2024; Salinas et al., 2023; Stuber et al, 2010; Xie et al, 2020)*. The discrepancy between our simulations and biosensor-based recordings versus those recording with FSCV measurements, is likely a combination of adsorption rates and the holding period between scans in voltammetry *(Burrell et al, 2015; Venton et al, 2002)*. To account for this, we inverted the deconvolution processes used in the literature to correct for DA kinetics observed with either voltammetry or amperometry (Extended Data Fig. 1C) *(Atcherley et al., 2015; Gonon et al, 2000; Venton et al., 2002)*. After this process, our results were very similar to those typically observed with FSCV (Fig. S1B) *(Atcherley et al., 2015; Brimblecombe et al., 2019; Stuber et al., 2010; Xie et al., 2020)*.

#### Supplementary Figures

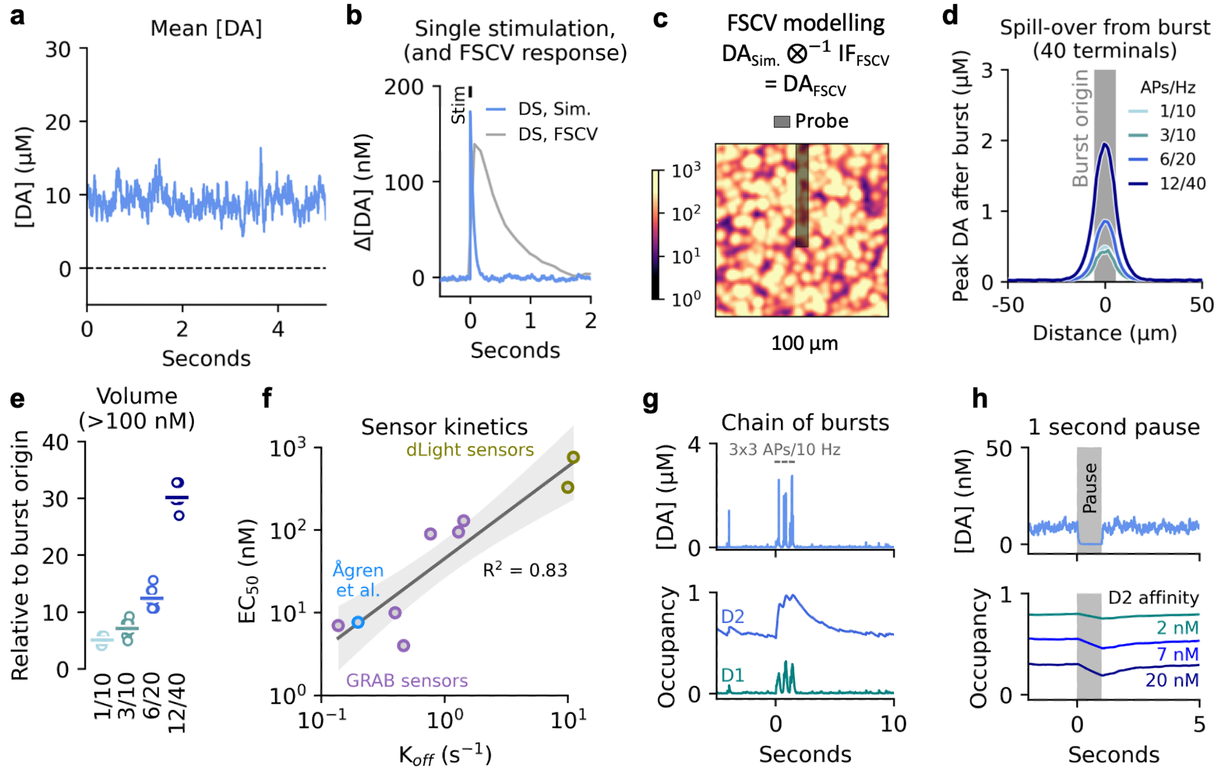


**Fig. S1.** Average concentration and receptor kinetics. (**a**) Average DA concentration across a 100x100x100 µm volume of simulated DS at pacemaker activity. (**b**) Mean concentration change in response to single event with 60% release probability (see supplementary notes on electrical stimulation) in all neurons at time zero. Blue trace is output directly from simulation, grey trace is the predicted FSCV measurement. (**c**) Modelling of predicted FSCV measurement and representative snapshot of simulation space just after a release event. (**d**) Peak DA concentration reached at different distances from area with phasic activity for the three firing scenarios. (**e**) Volume of space exposed to greater than 100 nM DA after firing relative to volume of space where terminals actively burst. (**f**) Least-square fit linear regression of the dLight and $GRAB_{DA}$ sensors based on reported kinetics *(Labouesse & Patriarchi, 2021)* and newest experimental characterization of D2R *(Agren et al, 2021)*. Shaded are indicates 95% C.I. (**g**) Top: representative [DA] trace for a voxel with a release site during pacemaker and a triple burst scenario (300 ms long bursts of 3 APs at 10 Hz, three times in a row with 300 ms in between). Bottom: Occupancy of D1Rs and D2Rs for same site. (**h**) Effect of complete pause in firing for 1 second on both average [DA] and D2R occupation at different affinities.

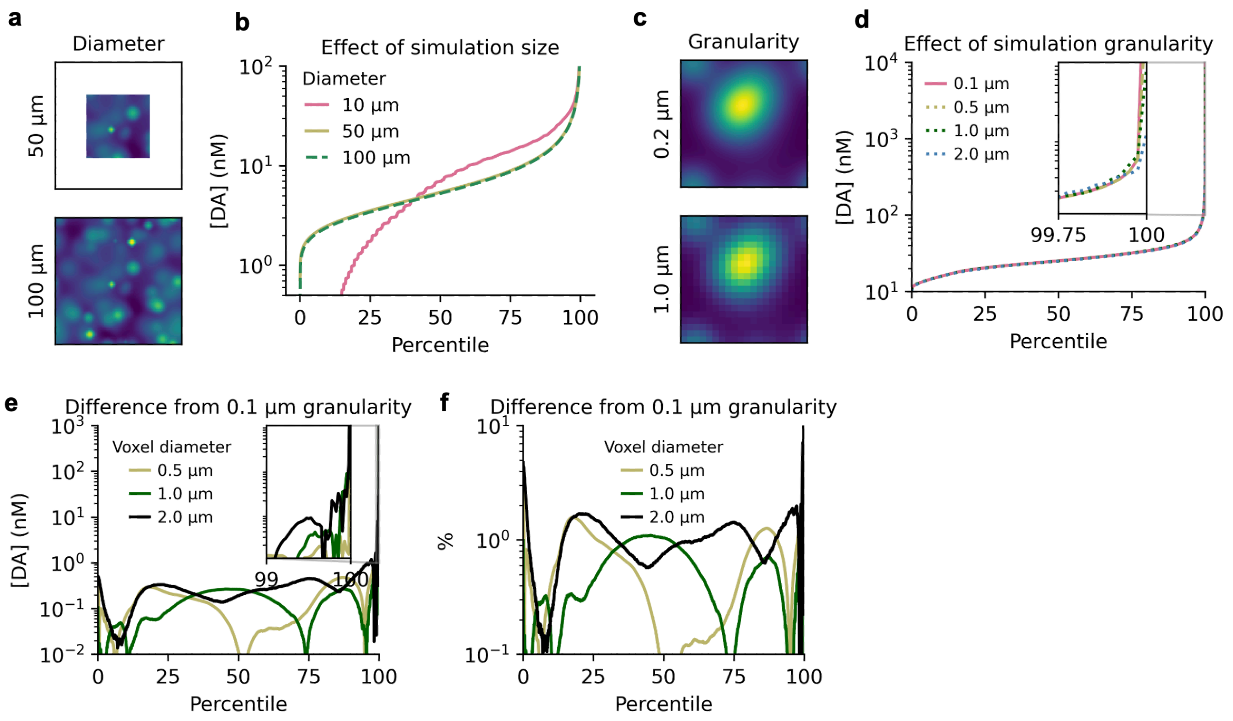


**Fig. S2.** Simulation size and granularity. (**a**) Schematic of different simulation sizes. (**b**) Concentration percentiles at different simulation diameters. Results are not robust until a diameter close to 50 µm is reached (line runs behind 100 µm line). (**c**) Schematic of simulation granularity. (**d**) Effect of simulation voxel diameter on concentration percentiles. Virtually no difference in concentration profiles below the 99.9$^{th}$ percentile of [DA], with the 1.0 µm voxel size still following 0.1 and 0.5 µm well above that level. Inset shows 99.75$^{th}$ to 100$^{th}$ percentile with y-axis matching main y-axis. (**e**) Absolute difference in [DA] between simulations at 0.1 µm voxel diameter and 0.5, 1.0 and 2.0 µm. Inset shows 99$^{th}$ to 100$^{th}$ percentile with y-axis matching main y-axis. (**f**) Percentage difference in [DA] between simulations at 0.1 µm voxel diameter and 0.5, 1.0 and 2.0 µm.

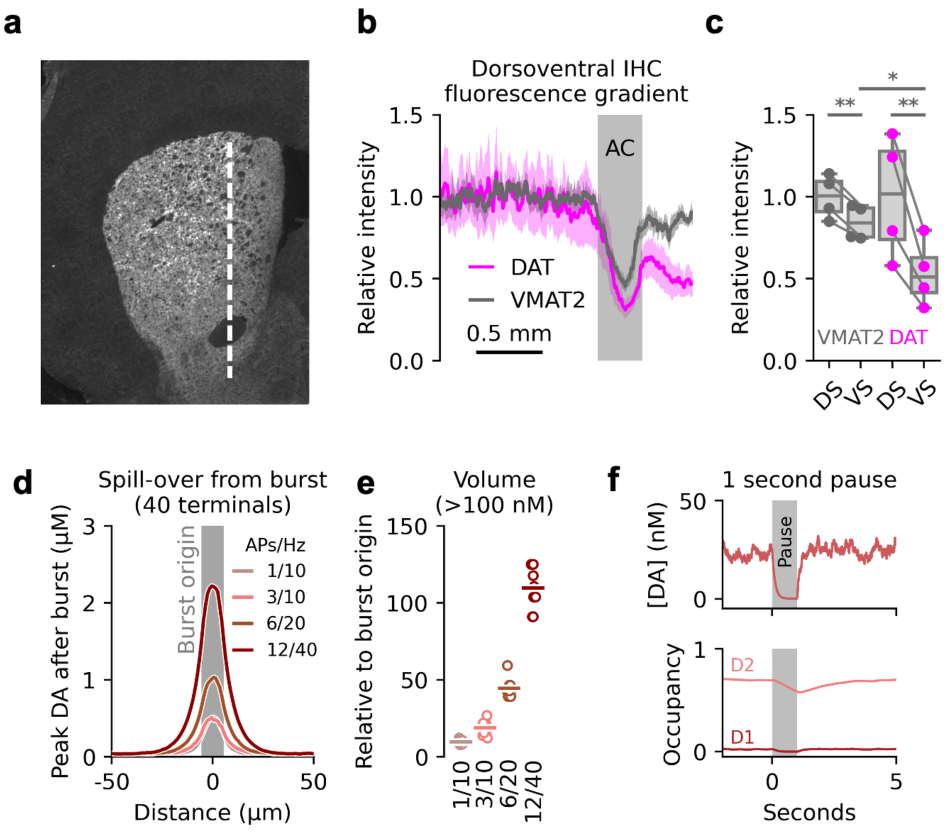


**Fig. S3.** Histochemical gradient of DAT and VMAT2 fluorescence. (**a**) Representative image of the mouse striatal slices analysed in b. Dashed white line indicates the quantified dorsoventral gradient. (**b**) Relative intensity of the DAT and VMAT2 immunosignal in the dorsoventral axis of striatal mouse brain slices from *(Sorensen et al, 2021)*. All slices show a drop at the anterior commissure (AC). Shaded areas around lines denote S.E.M. (**c**) Mean relative intensity of the DAT and VMAT2 signal before and after AC. Two-sided t-test, VS-DAT:VMAT2, P = 0.012(*), n = 4 mice; one-sided t-tests, DAT-DS:VS, P = 0.0021(**), VMAT2-DS:VS, P = 0.0086(**), n = 4 mice. (**d**) Peak DA concentration reached at different distances from area with phasic activity for the three firing scenarios in VS. (**e**) Volume of space in VS exposed to greater than 100 nM after firing relative to volume of space where terminals actively burst. (**f**) Effect of complete pause in firing in VS for 1 second on both average [DA] and D1R and D2R occupation.

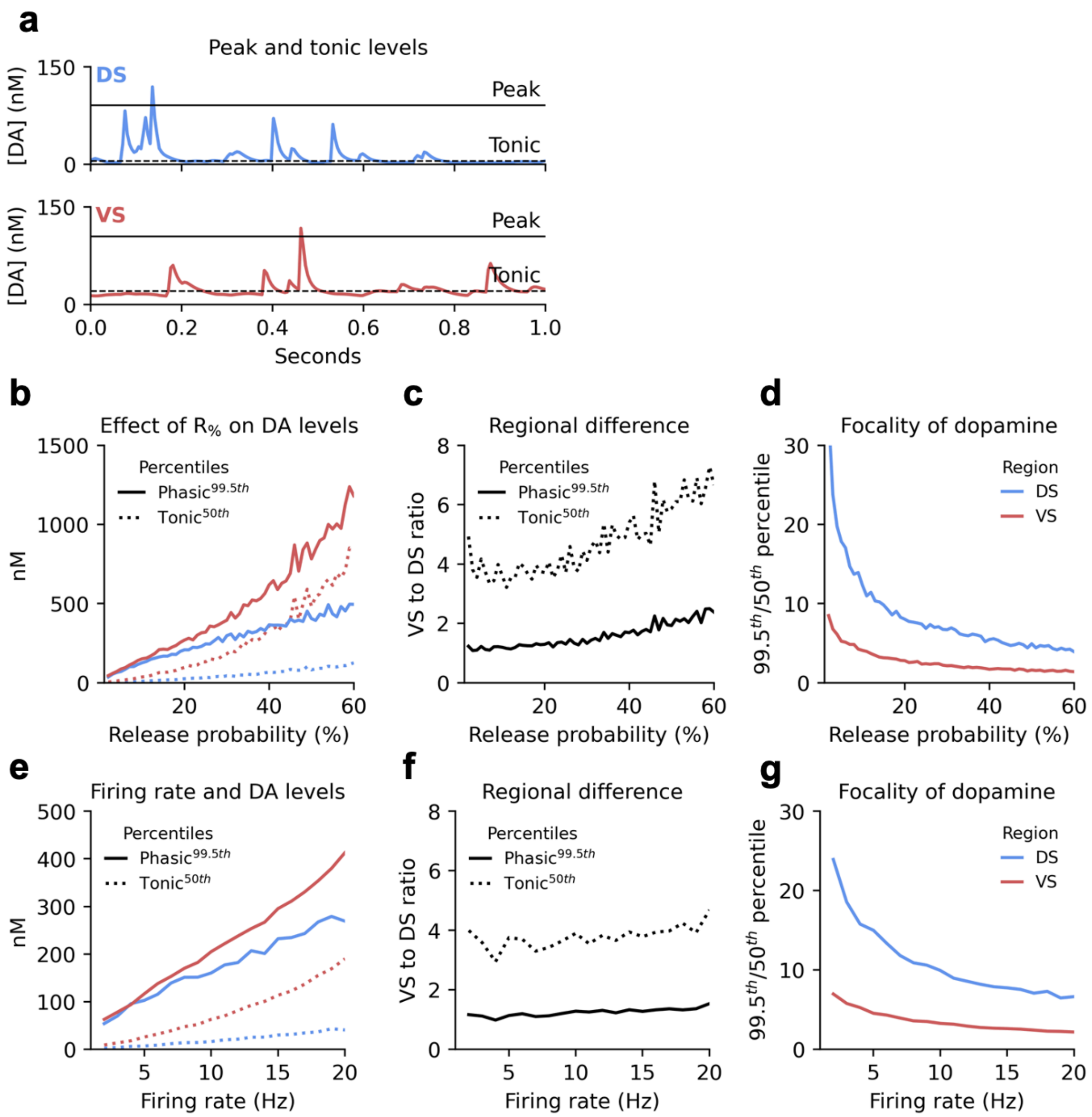


**Fig. S4.** Model parameter testing. (**a**) Schematic of our definitions of tonic (50$^{th}$ percentile/median, dashed lines) and peak (99.5$^{th}$ percentile, solid line) DA for both the dorsal and ventral striatum. (**b**) Effect of changing release probability ($R_{\%}$) on DA concentrations. (**c**) Relative difference between the ventral and dorsal striatum at different percentiles for different release probabilities. (**d**) Ratio between 99.5$^{th}$ and 50$^{th}$ percentile as a measure of focality for both regions. As $R_{\%}$ increases the concentrations becomes more homogeneous. (**e**) Effect of changing firing rate on DA concentrations. (**f**) Relative difference between the ventral and dorsal striatum at different percentiles for different firing rates. (**g**) Ratio between 99.5$^{th}$ and 50$^{th}$ percentile for both regions. As firing rate increases the concentrations becomes more homogeneous.

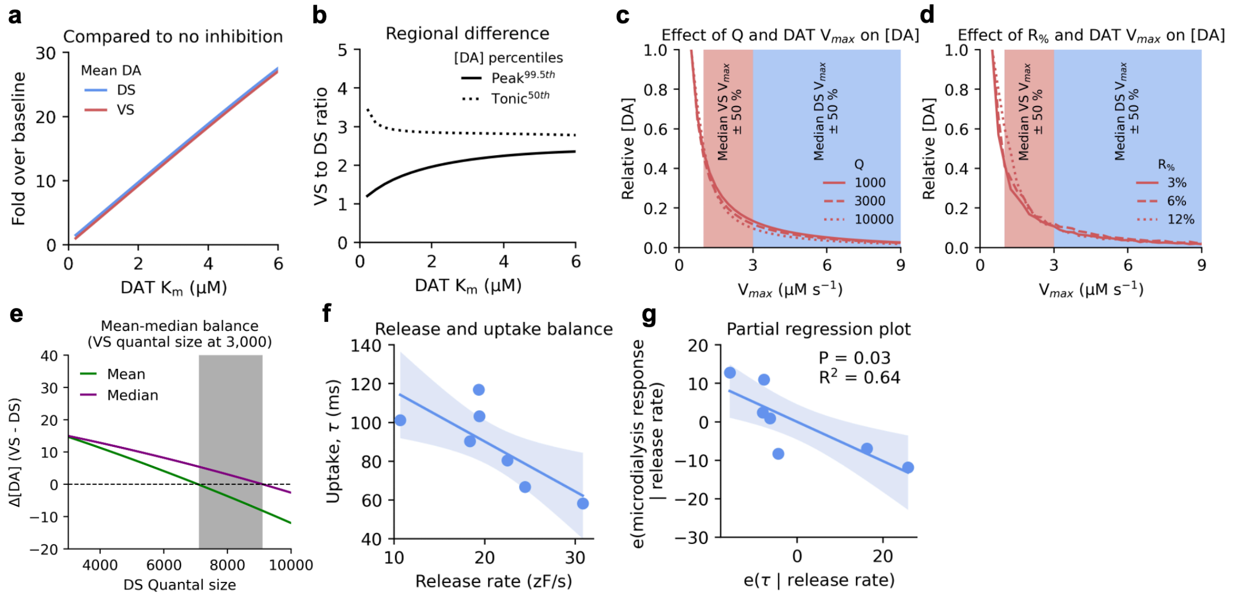


**Fig. S5.** Fold change during inhibition, $V_{max}$-sensitivity at different release parameters and release-uptake balance. (**a**) Fold change over baseline ($K_m$ of 210 nM) for mean DA concentration in the dorsal (DS) and ventral striatum (VS) with changing DAT $K_m$. (**b**) Relative difference between the dorsal and ventral striatum for both phasic and tonic DA at different $K_m$ values. (**c**) Effect of changing DAT $V_{max}$ on DA concentrations for three different quantal sizes (Q). [DA] normalized to highest values within each Q. Shaded area indicates median $V_{max}$ for DS and VS as found in the literature shown in Extended Data Table S2 with ± 50%. (**d**) Effect of changing DAT $V_{max}$ on DA concentrations for three different release probabilities ($R_{\%}$). [DA] normalized to highest values within each $R_{\%}$. Shaded area indicates median $V_{max}$ for DS and VS as found in the literature shown in Table S2 with ± 50%. (**e**) Difference between VS and DS for both mean and median concentrations across different Qs for DS (VS held at 3,000). Shaded area indicates range of Q where median is higher in VS but mean is higher in DS. (**f**) Least-square fit linear regression between release rate and autocorrelation decay rate ($\tau$) *(Ejdrup et al, 2023)*. Shaded area highlights 95% C.I. (**g**) Partial regression plot (error of the regression in (f) and error between DA response to amphetamine as measured by microdialysis and the release rate from *(Ejdrup et al., 2023)* to show that the less release and uptake correlate, the less release rate can explain the microdialysis response, suggesting release and uptake are partially independent of each other. Shaded area highlights 95% C.I.

#### Supplementary Tables

**Table S1.** Overview of reports on dopaminergic density and release in the striatum. Only studies that assessed both regions in rodents are included. Studies that stimulate directly in the striatum are omitted due to the large activation of nicotinic receptors on DA terminals (*1*, *2*). A.U. = arbitrary units, DS = dorsal striatum, VS = ventral striatum, TH = tyrosine hydroxylase.

| Measure | Ratio | DS | VS | Units | Species | Source |
|---|---|---|---|---|---|---|
| **TH expression density** | 100 % | ~90 | ~90 | A.U. | Mouse | *(Alberquilla et al, 2020)* |
| **TH immunoreactivity** | 95 % | ~68 | ~64 | A.U. | Mouse | *(Kuroda et al, 2010)* |
| **TH protein content** | 150 % | 0.07 | 0.11 | ng TH/µg prot. | Mouse | *(Salvatore et al, 2016)* |
| **DA content** | 90 % | ~155 | ~140 | ng DA/mg prot. | Mouse | *(Salvatore et al., 2016)* |
| **TH immunoreactivity** | 75 % | 2.8 | 2.1 | A.U. | Rat | *(Huang et al, 2019)* |
| **TH protein content** | 66 % | 0.36 | 0.24 | ng TH/µg prot. | Mouse | *(Salvatore et al, 2005)* |
| **FSCV - $[DA]_p$** | 91 % | 57 | 52 | nM | Rat | *(May & Wightman, 1989)* |
| **FSCV - $[DA]_p$** | 76 % | 89.3 | 67.5 | nM | Rat | *(Garris & Wightman, 1994)* |
| **Median** | 90 % | - | - | - | - | - |

**Table S2.** Overview of reported $V_{max}$ values for DA uptake in the striatum. Only studies that assessed both regions in rodents are included. DS = dorsal striatum, VS = ventral striatum, FSCV = fast-scan cyclic voltammetry.

| Method | VS/DS Ratio | DS (uM/s) | VS (uM/s) | Species | Source |
|---|---|---|---|---|---|
| **FSCV** | 29 % | 7.0 | 2.0 | Mouse | *(Calipari et al., 2012)* |
| **FSCV** | 28 % | 6.0 | 1.7 | Rat | *(Calipari et al., 2012)* |
| **FSCV** | 47 % | 3.0 | 1.4 | Rat | *(May & Wightman, 1989)* |
| **FSCV** | 31 % | 6.5 | 2.0 | Mouse | *(Siciliano et al., 2014)* |
| **FSCV** | 44 % | 5.0 | 2.2 | Rat | *(Ferris et al, 2014)* |
| **Median** | 31 % | 6.0 | 2.0 | - | - |
